# Mangroves deviate from other angiosperms in their genome size, leaf cell size, and cell packing density relationships

**DOI:** 10.1101/2022.09.12.507581

**Authors:** Guo-Feng Jiang, Su-Yuan Li, Russell Dinnage, Kun-Fang Cao, Kevin A. Simonin, Adam B. Roddy

## Abstract

**Background and Aims:** While genome size limits the minimum sizes and maximum numbers of cells that can be packed into a given leaf volume, mature cell sizes can be substantially larger than their meristematic precursors and vary in response to abiotic conditions. Mangroves are iconic examples of how abiotic conditions can influence the evolution of plant phenotypes.

**Methods:** Here, we examined the coordination between genome size, leaf cell sizes, and cell packing densities, and leaf size in 13 mangrove species across four sites. Four of these species occurred at more than one site, allowing us to test the effect of climate on leaf anatomy.

**Results:** We found that genome sizes of mangroves were very small compared to other angiosperms, and, like other angiosperms, mangrove cells were always larger than the minimum size defined by genome size. Increasing mean annual temperature of a growth site led to higher packing densities of veins (*D_v_*) and stomata (*D_s_*) and smaller epidermal cells but had no effect on stomatal size. Contrary to other angiosperms, mangroves exhibited (1) a negative relationship between guard cell size and genome size; (2) epidermal cells that were smaller than stomata, and (3) coordination between *D_v_* and *D_s_* that was not mediated by epidermal cell size. Furthermore, mangrove epidermal cell sizes and packing densities covaried with leaf size.

**Conclusions:** While mangroves exhibited coordination between veins and stomata and attained a maximum theoretical stomatal conductance similar to other angiosperms, the tissue-level tradeoffs underlying these similar relationships across species and environments was markedly different, perhaps indicative of the unique structural and physiological adaptations of mangroves to their stressful environments.

## INTRODUCTION

One of the more iconic examples of how the environment can select for plant phenotypes are mangroves. The mangrove habitat is characterized by multiple stresses, including osmotic and drought stress, tidal inundation, high winds, high temperature, and high ultraviolet (UV) radiation, which influence gas exchange rates of leaves and plant survival (Tomlinson, 1986; Ball, 1988; Krauss *et al*., 2008). Together, these abiotic conditions make the mangrove habitat particularly stressful and suggest that mangroves are a valuable resource for understanding plant adaptation to extreme environments (Scholander *et al*., 1964; Reef and Lovelock, 2015). The mangrove habit has evolved repeatedly in over 20 lineages of vascular plants (Duke, 1992; He *et al*., 2022) and encompasses both convergent evolution of similar traits as well as the evolution of multiple novel morphological, anatomical, and physiological strategies to survive under similar environmental challenges. These adaptations include diverse leaf morphologies (e.g., leaves with glands that secrete salt), extensive support roots, buttress roots, and viviparous water-dispersed propagules (Tomlinson, 1986; Ball, 1988; Reef and Lovelock, 2015). The mangrove habitat is often considered extreme, with warm temperatures characteristic of the tropics, frequent wind characteristic of coastal shores, and near constant saltwater inundation from the ocean.

Because growth and survival depend on maintaining physiological function in the face of often stressful environmental conditions, leaf anatomical traits that influence rates of photosynthetic carbon gain may show plastic responses to the environment within species and vary among species associated with their habitat affinities. Of particular importance are the leaf anatomical traits that limit diffusion of CO_2_ across the leaf surface and into the mesophyll cells where photosynthesis occurs (Franks and Beerling, 2009; Brodribb *et al*., 2010; Boyce and Zwieniecki, 2012; Franks *et al*., 2012a; Franks *et al*., 2012b; Théroux-Rancourt *et al*., 2021). Leaf surface conductance and the anatomical traits that control maximum potential CO_2_ diffusion into the leaf act as first-order constraints on photosynthetic capacity and, therefore, define an upper limit to how much carbon can be allocated to growth, reproduction, and defense (Franks and Beerling, 2009; Roddy *et al*., 2020). Increasing leaf surface conductance to CO_2_ has occurred predominantly by decreasing the size (*S_s_*) and increasing the packing density (*D_s_*) of stomata (Franks and Beerling, 2009). Opening stomata to allow CO_2_ diffusion into the leaf exposes the wet surfaces of the leaf mesophyll cells to a dry atmosphere, resulting in evaporative water loss that must ultimately be replaced in order to maintain water balance and physiological function. As a result, increasing *D_s_* is generally associated with a higher density of leaf veins (*D_v_*), which efficiently supply liquid water throughout the leaf (Sack and Frole, 2006; Brodribb *et al*., 2007; Boyce *et al*., 2009; Brodribb, 2009; de Boer *et al*., 2012).

Coordination between veins and stomata is thought to be critical to maintaining leaf water balance, and correlations between *D_s_* and *D_v_* suggest coordinated development across multiple cell types and tissues during development. Because more cells and cell types can be packed into a given space if these cells are smaller, reducing cell size has been a primary way of increasing the packing densities of multiple tissue types, including stomata, veins, and mesophyll cells (Franks and Beerling, 2009; Brodribb *et al*., 2013; Simonin and Roddy, 2018; Théroux-Rancourt *et al*., 2021). How small a cell can be (i.e. meristematic cell size)– and, by extension, the maximum number of cells that can be packed into a given space–is fundamentally limited by the volume of the nucleus, or, as is more commonly measured, genome size (Cavalier-Smith, 1978; Beaulieu *et al*., 2008; Šímová and Herben, 2012; Simonin and Roddy, 2018; Roddy *et al*., 2020). Smaller cells and mesophyll tissues composed of smaller cells have higher surface area-to-volume ratios, which allow for higher rates of CO_2_ diffusion into photosynthetic tissues (Théroux-Rancourt *et al*., 2021). While genome downsizing, particularly among angiosperm lineages during the Cretaceous, was critical in reducing minimum cell sizes and allowing for smaller, more densely packed cells (Simonin and Roddy, 2018), genome downsizing also allows for a greater range of mature cell sizes and packing densities (Roddy *et al*., 2020; Théroux-Rancourt *et al*., 2021). Mature cells are often considerably larger than their meristematic precursors, and the process of cell expansion allows cell sizes and packing densities to be tuned to environmental conditions. For example, differential expansion of epidermal pavement cells during leaf development can lead to coordinated changes in the densities of veins and stomata (Carins Murphy *et al*., 2012, 2014, 2016, 2017). Thus, while minimum cell sizes and maximum cell packing densities are limited by genome size, deviation from these extreme limits may be driven by differential expansion of leaf cells in response to variable environmental conditions (i.e. trait plasticity). Thus, abiotic factors that influence plant water balance, and by extension of cell expansion, can cause deviation in leaf anatomical traits away from the extreme limits imposed by genome size.

Here we sought to characterize (1) how genome size limits cell sizes and packing densities in mangrove leaves, (2) how abiotic conditions influence intraspecific variation in anatomical traits, and (3) how traits of different cell types that influence leaf function are coordinated within and among mangroves compared to other non-mangrove angiosperms. We sampled a total of 13 species (one of them is a naturally occurring hybrid) from four sites (Fig. 1), with four of these species occurring at more than one site. We explicitly incorporated previously published data for comparison to mangroves and included phylogenetically corrected regressions. Our results showcase that while some scaling relationships defined for angiosperms apply to mangroves as well, mangroves nonetheless deviate in other relationships, and these deviations may be due to the unique conditions of the mangrove habitat. Understanding leaf trait coordination in hardy plants like mangroves provides an important test of current theory linking leaf structure to leaf function.

**Fig. 1.**
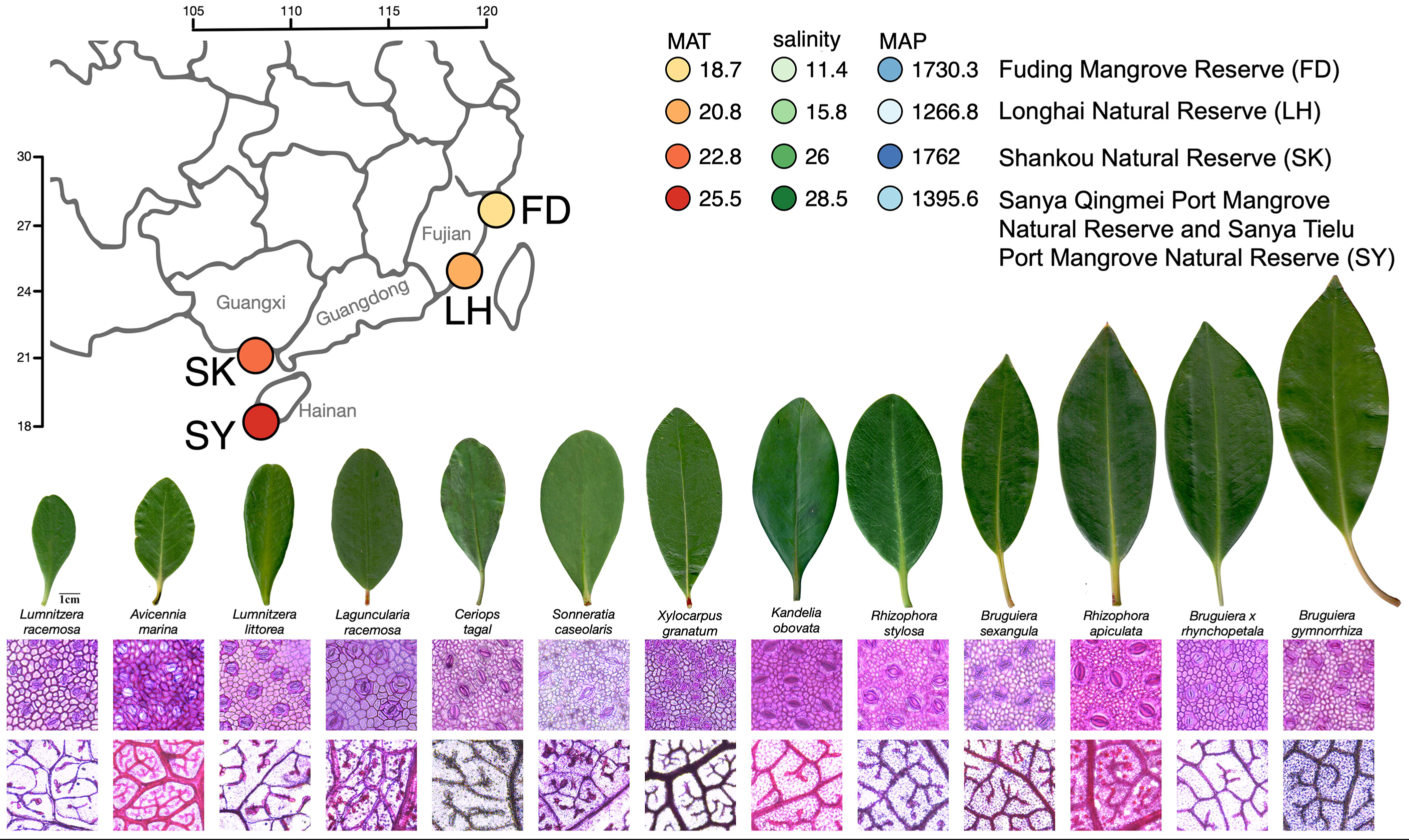
Sites and species sampled. Map of southeastern China shows the four sites in the provinces of Fujian, Guangxi, and Hainan where plants were sampled. Points on the map are colored according to their mean annual temperature (MAT), and colors for site means of annual temperature (MAT, °C), soil water salinity (g kg^−1^), and annual precipitation (MAP, mm), are shown to the right. Note these colors are used in subsequent figures to color points. Images of exemplary leaves of each species show the diversity in leaf size among the species. Scale bar next to the leaf of *Lumnitzera racemosa* is 1 cm and applies to all leaf images. Microscopic images below each leaf are exemplary images of abaxial epidermises for each species, and each epidermal image is 250 μm on each side. The bottom row of images shows exemplary images of veins for each species, and each vein image is 1000 μm on each side.

## MATERIALS AND METHODS

### Study sites and plant material

Mangrove plants were sampled in five natural reserves along a latitudinal gradient in Southern China (Fig. 1; Table 1): Fuding Mangrove Natural Reserve (FD; 27° 20’ N / 120° 12’ E), Longhai Mangrove Natural Reserve (LH; 24° 29’ N / 118° 04’ E), Shankou Mangrove Natural Reserve (SK; 21° 28’ N/109° 43’ E), Sanya Tielu Port Mangrove Natural Reserve (SYTL; 18° 15’ N / 109° 42’ E), and Sanya Qingmei Port Mangrove Natural Reserve (SYQM; 18° 14’ N / 109° 36’ E), the latter two are located about 7 km from each other and so were grouped as the same site Sanya (SY) in our analysis. In total, 13 species were collected across all sites, and all taxa except *Kandelia obovata* occurred at the southernmost site SY (Table 2). *K. obovata* was the only species that occurred at the northernmost site (FD). Three species were found in three sites, and another species occurred in two sites (Table 2). These four species located at multiple sites were used to test for environmental effects on anatomical traits.

**Table 1.** Site names, locations, and mean climate variables used in the analyses.

| *Site name* | *Latitude, longitude* | *Mean annual temperature (MAT) from years 1955-2016 (°C)* | *Mean Annual precipitation (MAP) from years 1955-2016 (mm)* | *Mean relative humidity (%)* | *Salinity (g/kg)* |
| --- | --- | --- | --- | --- | --- |
| Fuding Mangrove Natural Reserve (FD) | 27° 20' N, 120° 12' E | 18.7 | 1730.3 | 78.1 | 11.4 |
| Longhai Mangrove Natural Reserve (LH) | 24° 29' N, 118° 04' E | 20.8 | 1266.8 | 77.0 | 15.8 |
| Shankou Mangrove Natural Reserve (SK) | 21° 28' N, 109° 43' E | 22.8 | 1762.0 | 80.4 | 26.0 |
| Sanya Tielu Port Mangrove Natural Reserve and Sanya Qingmei Port Mangrove Natural Reserve (SY) | 18° 15' N, 109° 42' E<br>18° 14' N, 109° 36' E | 25.5 | 1395.6 | 79.2 | 28.5 |

**Table 2.** Taxa sampled, their taxonomic authorities, and the sites at which they occurred. Location abbreviations are as follows: FD, Fuding Mangrove Natural Reserve; LH, Longhai Mangrove Natural Reserve; SK, Shankou Mangrove Natural Reserve; SYTL, Sanya Tielu Port Mangrove Natural Reserve; SYQM, Sanya Qingmei Port Mangrove Natural Reserve.

| *Family* | *Species* | *Location* |
| --- | --- | --- |
| Rhizophoraceae | *Kandelia obovata* (S., L.) Yong | FD |
| Rhizophoraceae | *Kandelia obovata* (S., L.) Yong | LH |
| Rhizophoraceae | *Bruguiera gymnorrhiza* (Linn.) Savigny | LH |
| Acanthaceae | *Avicennia marina* (Forsk.) Vierh. | LH |
| Rhizophoraceae | *Kandelia obovata* (S., L.) Yong | SK |
| Rhizophoraceae | *Bruguiera gymnorrhiza* (Linn.) Savigny | SK |
| Rhizophoraceae | *Rhizophora stylosa* Griff. | SK |
| Acanthaceae | *Avicennia marina* (Forsk.) Vierh. | SK |
| Rhizophoraceae | *Rhizophora apiculata* Bl. | SYTL |
| Rhizophoraceae | *Bruguiera sexangula* (Lour.) Poir. | SYTL |
| Rhizophoraceae | *Bruguiera x rhynchopetal* W. C. Ko. | SYTL |
| Rhizophoraceae | *Bruguiera gymnorrhiza* (Linn.) Savigny | SYTL |
| Meliaceae | *Xylocarpus granatum* Koenig. | SYTL |
| Combretaceae | *Lumnitzera racemosa* Willd. | SYTL |
| Combretaceae | *Lumnitzera littorea* (Jack) Voigt | SYTL |
| Combretaceae | *Laguncularia racemosa* (L.) C.F.Gaertn. | SYTL |
| Acanthaceae | *Avicennia marina* (Forsk.) Vierh. | SYTL |
| Sonneratiaceae | *Sonneratia caseolaris* (L.) Engl. | SYTL |
| Rhizophoraceae | *Ceriops tagal* (perr.) C. B. Rob. | SYQM |
| Rhizophoraceae | *Rhizophora stylosa* Griff. | SYQM |

At least three randomly selected individuals per species per site were selected for sampling. Sun-exposed branches were cut and sealed in a plastic bag with wet tissues, then transported back to the laboratory at Guangxi University for subsequent sample processing and measurements.

### Anatomical traits

All measurements were made on three to six randomly selected, fully expanded, healthy, sun-exposed leaves of each species at each site. Three to six approximately 1-cm^2^ sections of lamina were sampled from each leaf, avoiding the leaf margin and midrib. These sections were cleared in a 1:1 solution of 30% H_2_O_2_ and 100% CH_3_COOH and incubated at 70 °C until all pigments had been removed. The sections were then rinsed in water and the epidermises separated with forceps from the mesophyll and veins, allowing these three layers (upper epidermis, lower epidermis, and mesophyll with veins) to be stained and mounted separately. To increase contrast, all samples were stained with Safranin O (1% w/v in water) for 15 min and Alcian Blue (1 % w/v in 3 % acetic acid) for 1 minute, then washed in water and mounted on microscope slides. We found that Safranin O and Alcian Blue did not both readily bind to the mangrove leaves.

Images were taken at 5x, 10x, or 20x magnification, which had fields of view of approximately 3.99 mm^2^, 0.89 mm^2^, and 0.22 mm^2^, respectively, using a compound microscope outfitted with a digital camera (DM3000, Leica Inc., Germany). Both abaxial (lower) and adaxial (upper) leaf surfaces were imaged for all species because *Laguncularia*, *Lumnitzera*, and *Sonneratia* species were known to have stomata on both surfaces. In the following analysis, we used only abaxial (bottom surface) *D_s_* for packing densities, and the sum of adaxial and abaxial stomatal densities (termed *D_s,tot_*) for analyses related to fluxes.

All anatomical measurements from images were made using ImageJ (Rueden *et al*., 2017). From images of paradermal sections, vein density (*D_v_*) was measured as the total length of leaf vascular tissue per mm^2^ of leaf area, epidermal pavement cell size (*S_ec_*) was quantified by measuring the two-dimensional area of individual epidermal cells in an epidermal image, guard cell length (*l_g_*) was measured as the maximum length of one guard cell in a pair, stomatal density (*D_s_*) was measured by counting the number of stomata in the image and dividing by the area of the field of view, and epidermal cell packing density (*D_ec_*) was measured by counting the number of epidermal cells in an image and dividing by the area of the field of view. Partial stomata and epidermal cells were included in the density counts if they were partially bisected by the top and left borders of the image and ignored if they were partially bisected by the bottom and right borders of the image. *S_ec_* was measured on approximately ten randomly chosen epidermal cells that were not touching stomata in each image. Measurements of *l_g_* were made on ten stomata per image.

We compared two methods for estimating the two-dimensional projected surface area of stomata (i.e. stomatal size, *S_s_*) in the plane of the leaf epidermis. First, we manually measured the area of each of 10 stomata per image. Second, we calculated the two-dimensional area of one guard cell as:

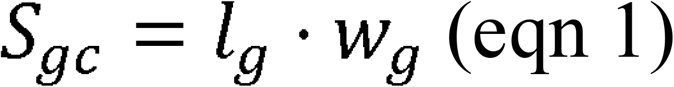

where *l_g_* is the length of one guard cell and *w_g_* is the width of one guard cell, which can be estimated as w_g_ = l_g_ · 0.36 (de Boer *et al*., 2016). Doubling *S_gc_* is equivalent to the size of a pair of guard cells, i.e. one stoma (*S_s_*). Species’ average estimates of *S_s_* using these two methods were strongly correlated (*R^2^* = 0.90, P < 0.0001) with a standard major axis slope not significantly different from unity (P = 0.11), and so for subsequent analyses we used the measured values of *S_s_*.

Epidermal cell size (*S_ec_*) was quantified in two ways. First, the average *S_ec_* for an image was calculated according to Carins-Murphy et al. (2017) as:

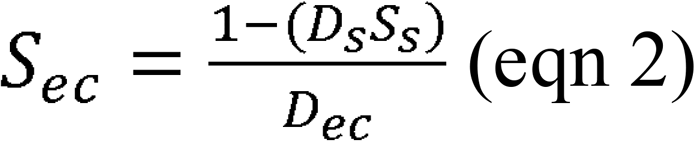

where *D_s_* is stomatal density, *S_s_* is stomatal size, and *D_ec_* is the epidermal cell density. Second, we directly measured the two-dimensional surface area of five randomly chosen epidermal cells in each image. These two methods showed strong agreement in quantifying the two-dimensional epidermal cell surface area (R^2^ = 0.86, P < 0.0001) with a slope not significantly different from unity (P = 0.93), and so for subsequent analyses we used the direct measurements of pavement cell surface area.

### Genome size

The genome sizes of the mangrove species studied here were taken from the literature (Lyu *et al*., 2018; Hu *et al*., 2020; He *et al*., 2022). Measurements of genome size in megabases (Mb) were converted to picograms (pg) following the equation 1 pg = 1 Mb / 978 (Dolezel *et al*., 2003.

### Environmental data for the four sampling sites

Climate data (mean annual temperature, MAT, and mean annual precipitation, MAP) for each site were downloaded from http://data.cma.cn. Climate data were long-term averages of monthly collected raw data from January 1951-December 2016. Estimates of soil water salinity were obtained from published references for FD (Lin *et al*., 1998) and LH (Cao, 2008), while data for SK and SY were provided by the Guangxi Mangrove Research Center, Guangxi Academy of Sciences (previously unpublished data). At all sites, soil samples of 0-50 cm depth were sampled throughout the area where the plants were growing, from inland to the seashore, during low tide. At least 5 soil samples were collected for each site with PVC tubes, then the soils were stored in plastic bags, and transported back to the laboratory. Soil water was collected by filtering the soil from the water (Cao, 2008).

### Modeling gas exchange capacity from anatomical traits

Using these anatomical data, we modeled maximum (*g_s,max_*) stomatal conductance using previously published methods. Maximum theoretical *g_s_* was calculated according to Franks and Beerling (2009) as:

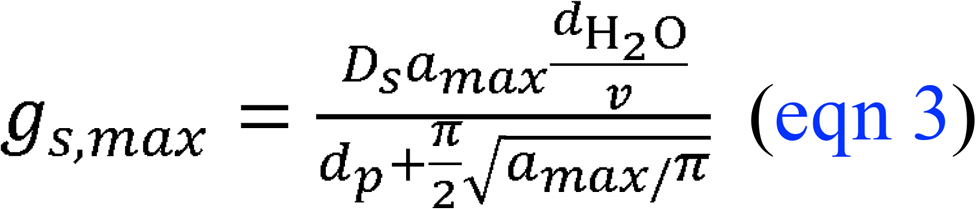

where *d*_H2O_ is the diffusivity of water vapor in air (0.0000249 m^2^ s^−1^), *v* is the molar volume of air normalized to 25°C (0.0224 m^3^ mol^−1^), *D_s_* is the stomatal density, *d_p_* is the depth of the stomatal pore, and *a_max_* is the maximum area of the open stomatal pore. The depth of the stomatal pore, *d_p_*, was assumed to be equal to the width of one guard cell, which was taken as 0.36 ⋅ *l_g_* (de Boer *et al*., 2016). The maximum area of the open stomatal pore, *a_max_*, was approximated as π(*p*/2)^2^ where *p* is stomatal pore length and is approximated as *l_g_*/2. Thus, *g_s,max_* could be calculated from measurements of *l_g_* and *D_s_*.

### Leaf mass per area (LMA) measurements

For LMA determination, 15 leaves were randomly selected per species, and their areas were measured with an LI-3000A (LI-COR, USA). Afterwards, samples were oven-dried at 70°C for 72 hours, weighed, and LMA was calculated as leaf dry mass divided by fresh leaf area. Leaf size values were obtained from these leaf area data.

### Previously published data

To generate a broad, phylogenetically diverse dataset of angiosperm leaf anatomical traits, we compiled data from Beaulieu et al. (2008), Blonder and Enquist (2014), Boyce et al. (2009), Brodribb and Feild (2010), Brodribb et al. (2013), Jordan et al. (2013), Coomes et al. (2008), Feild et al. (Feild *et al*., 2009; Feild *et al*., 2011), Fridley and Craddock (2015), Gleason et al. (2016), Sack et al. (2012), Carins Murphy et al. (2016), Bongers and Popma (1990), and McElwain et al. (2016). Taxonomic names were corrected by comparing querying The Plant List using the R package “Taxonstand” (Cayuela *et al*., 2012). We merged these data with the Kew Plant DNA C-Values database (version 6), after using the same procedure to check taxonomic names in that database. Because our focus was on plants with capsule-shaped guard cells, we removed monocots from the dataset because they are known to have dumbbell-shaped guard cells. The resulting database encompassed 836 species from 126 families. Of these species, there were 300 species from 68 families with guard cell length data, 274 species from 62 families with stomatal density data, and 638 species from 111 families with vein density data. There were 289 species with both *l_g_* and *D_s_* measurements. Meristematic cell volumes as a function of genome size were taken from Šímová and Herben (2012). Using these measured volumes of meristematic cells, we approximated the maximum two-dimensional cross-sectional area of a spherical meristematic cell by calculating the cross-sectional area of a sphere with the same volume.

### Data analysis

All statistical analyses were conducted in R (v. 4.0.3) (R Core Team 2020). We used linear regression and standard major axis (SMA) regression (R package ‘smatr’) to determine the relationships between traits (Warton *et al*., 2012). SMA regressions were used on log-scaled traits. For visualization, confidence intervals around SMA regressions were calculated from bootstrapping the SMA regressions 1000 times. We used slope tests, implemented in ‘smatr’ to compare slopes and report P-values for whether the slopes are significantly different or not. For the relationships between genome size and the sizes of guard cells and epidermal pavement cells, we calculated the SMA regressions and associated statistics using species means (i.e. averaged across sites) because species occurred at different numbers of sites, but we plot points representing the species x site means to better visualize the variation in cell size within species across sites. Because we did not measure leaf size and LMA on the same leaves on which anatomical measurements were made, analyses of leaf size use species x site mean data for each variable and rely on linear regressions.

To determine the effects of climate variables on anatomical and physiological traits, we constructed linear mixed effects models for each trait and climate variable, using the four species that were present at more than one site and including measurements from individual plants (i.e. not using species x site mean trait values). We used the R package ‘lme4’ to construct models that had a fixed effect of the environmental variable and a random effect of species (Bates *et al*., 2014). This random effect allowed each species to have a different intercept. Because ‘lme4’ is unable to incorporate uncertainty about the random effects into predictions, confidence intervals around the fixed effects cannot be calculated. We report two *R^2^* values: (1) the marginal *R^2^* and P-value of the environmental variable after accounting for the differences among species (i.e. random effects), and (2) the conditional *R^2^* that indicates how much variation is explained by the entire model (i.e. both the environmental variable and species identity).

To account for the statistical non-independence of sampling related species, we incorporated phylogenetic covariance into our regression analyses. For mangroves, we used a recently published chloroplast phylogeny that included the species sampled here (Li *et al*. 2021a; Li *et al*. 2021b), and we used trait data for the southernmost population of each species (i.e. Sanya for all species except *Kandelia obovata*, for which the southernmost population was located at Shankou Natural Reserve). For non-mangrove angiosperms, we constructed a broad phylogeny based on the large dated seed plant phylogeny of Smith and Brown (2018), hereafter the ‘reference phylogeny’. Approximately 80% of our species were included in the reference phylogeny. We placed the remaining 47 species onto the reference phylogeny using a randomization procedure based on known taxonomic relationships. Forty-five of the 47 species were placed within a congeneric clade. The remaining two species were placed with other members of the same tribe (*Pseudolmedia glabrata* within the Castilleae and *Trichospermum mexicanum* within the Grewieae). Rather than just placing taxa randomly (i.e. uniformly) within the clade, the procedure for placement attempted to preserve the relative distribution of branch-lengths within different clades, which is becoming the standard practice (Chang *et al*., 2020, Thomas *et al*., 2013). Rather than resolving the branch-lengths using a fitted diversification model, such as a birth-death model (as in Chang *et al*., 2020), we used a non-parametric approach with the goal of preserving clade-level branch-length distributions. Based on the reasoning that the tips found in a clade on the reference tree represent a sample drawn from the true diversification history of a clade, we placed missing species by replacing randomly chosen species from the target clade on the reference phylogeny with the target missing species. This is conceptually a bootstrapping approach to missing species placement. The one exception to this approach were 7 species in the genus Trimenia, for which the reference phylogeny had only one member. Since we had no other phylogenetic data for this genus, these species were placed in a polytomy with zero branch-lengths between them. After all missing species were placed, we pruned the reference tree of all species that were not in our dataset. We created a distribution of trees representing phylogenetic uncertainty by repeating this process 1000 times, generating 1000 equally likely alternative placements for missing species.

For mangroves and non-mangrove angiosperms we used phylogenetically corrected regressions between pairwise trait combinations using the R packages ‘ape’ (Paradis and Schliep, 2019) and ‘picante’ (Kembel *et al*., 2010). We calculated phylogenetically corrected generalized least squares regressions for pairwise trait combinations using the R function *gls* with a correlation structure equivalent to the phylogenetic relatedness under a Brownian motion model of evolution (R function *corBrownian*). In order to account for the phylogenetic uncertainty of the non-mangrove angiosperms in the previously published data, we calculated these phylogenetic regressions on all 1000 equally likely phylogenies and report the distributions of test statistics (slope, *t*-statistics, P-value) in the supplemental information.

## RESULTS

### Relationships between genome size and cell size in mangroves and non-mangrove angiosperms

Among non-mangrove angiosperms, there was a significant, positive relationship between guard cell size (*S_gc_*, μm^2^) and genome size (Fig. 2; slope = 0.50 [0.44, 0.56], R^2^ = 0.62, P < 0.0001) that remained highly significant after accounting for shared phylogenetic history (Figure S1). However, among mangroves the relationship between guard cell size (*S_gc_*, μm^2^) and genome size was negative (Fig. 2; slope = −1.16 [−1.81, −0.74], R^2^ = 0.34, P = 0.02), such that species with larger genomes had smaller guard cells (Fig. 2), but this relationship was not significant after accounting for shared evolutionary history (P = 0.27). Moreover, there was no effect of genome size on epidermal pavement cell size among mangroves (*S_ec_*, μm^2^; non-phylogenetic P = 0.37; phylogenetic P = 0.08), although among a broader sampling of angiosperms epidermal cell size and genome size were strongly and positively correlated (slope = 0.50 [0.44, 0.56], R^2^ = 0.59, P < 0.0001) even after accounting for shared evolutionary history (Figure S1). Additionally, *S_gc_* was generally greater than *S_ec_* among mangroves (*S_gc_* / *S_ec_* = 2.91 ± 0.55, compared to *S_gc_* / *S_ec_* = 0.18 ± 0.02 among non-mangrove angiosperms) and both cell types were always larger than the minimum cell size defined by the sizes of the genome (Fig. 2).

**Fig. 2.**
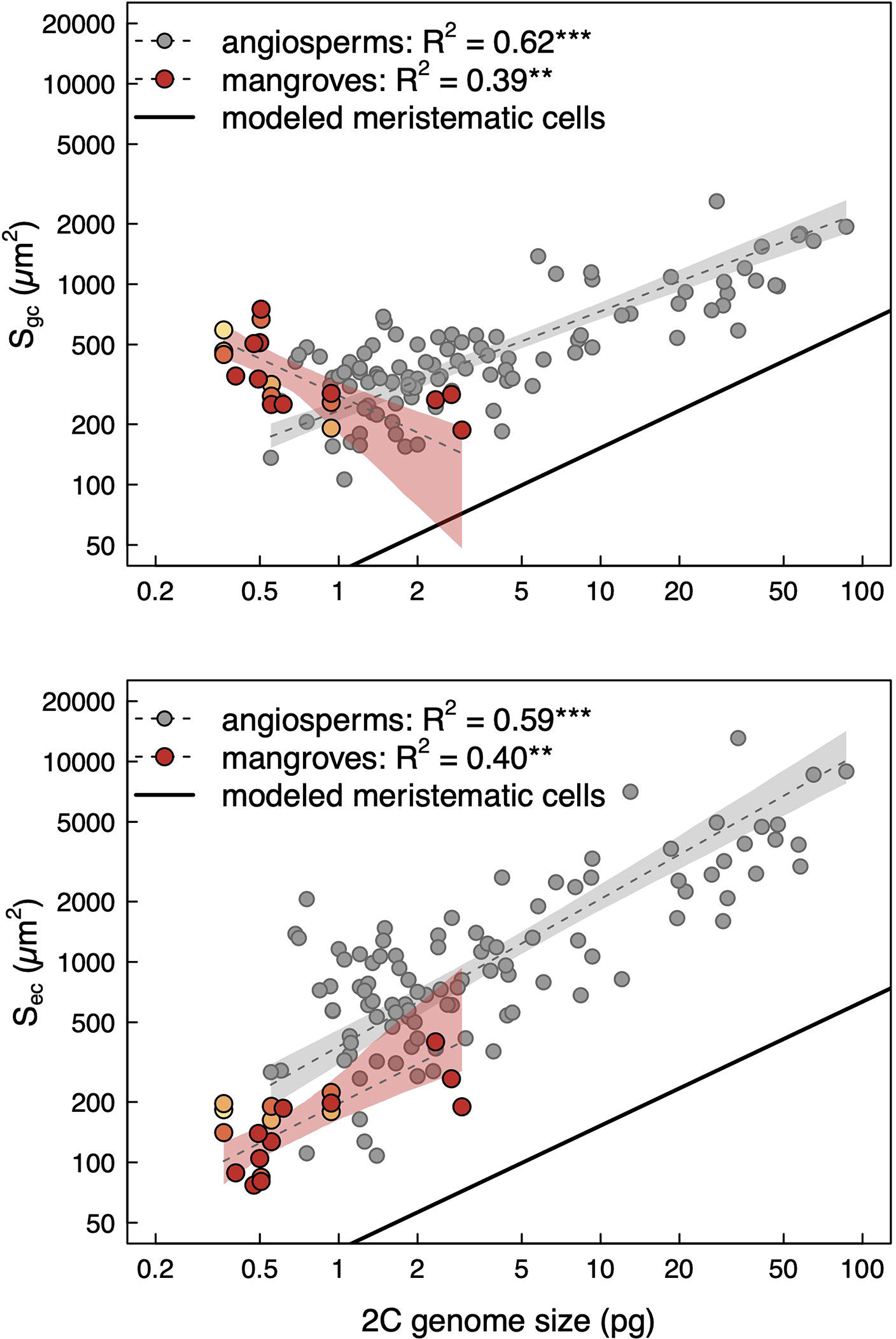
Two-dimensional sizes of (a) stomatal guard cells (*S_gc_*) and (b) epidermal pavement cells (*S_ec_*) as a function of genome size. The solid line indicates the maximum two-dimensional cross-sectional area of meristematic cells based on data from Šímová and Herben (2012) (see methods for details about calculation). *S_gc_* was calculated from measurements of guard cell length assuming that guard cells were shaped as capsules (see methods for details). *S_ec_* was measured from epidermal images. Note that points for mangroves represent species x site means and standard errors, though standard error bars are small enough that they are mostly obscured by the points. Grey points represent published data from angiosperms.

### Relationships between mean annual temperature and leaf anatomical traits

The final sizes of mature cells are often much larger than their meristematic precursors, allowing final cell size to be influenced by environmental variation. For the four species that occurred at multiple sites, we tested the effects of climate on leaf anatomical traits by incorporating intraspecific variation in traits and a random effect of species (Fig. 3). The conditional *R^2^* that indicates the amount of variation explained by the entire model (i.e. the fixed effect of the environmental variable and the random effect of species) were all above 0.50 and often above 0.75, indicating that the there was substantial variation among species in how traits responded to environmental variation (Figs. 3, S2; Table S1). The marginal *R^2^*, which indicates the amount of variation explained by the environmental variable alone, were much lower though often still significant. Increasing temperature (MAT) had significant effects on the packing densities of stomata (marginal R^2^ = 0.07, P < 0.01) and leaf veins (marginal R^2^ = 0.03, P < 0.01) and the sizes of epidermal cells (marginal R^2^ = 0.03, P < 0.01), but not on the sizes of stomata (*S_s_*) or the packing densities of epidermal cells (*D_ec_*) (Fig. 3). The small marginal R^2^ combined with high conditional R^2^ reported here indicate that climate did have a significant but very small effect on traits, but species identity had a much larger effect on the traits. In other words, each species was very different even at the same site, and the species responded similarly but by a small amount to the environmental differences across sites. There were no significant effects of MAP on any of the anatomical traits (all P > 0.05; Fig. S2, Table S1). Because MAT and salinity were strongly correlated across sites (Fig. 1), higher salinity significantly increased the packing densities of stomata (marginal R^2^ = 0.08, P < 0.001) and leaf veins (marginal R^2^ = 0.03, P = 0.001), but salinity had no significant effect on the sizes of stomata or epidermal cells or on the packing densities of epidermal cells (Fig. S2).

**Fig. 3.**
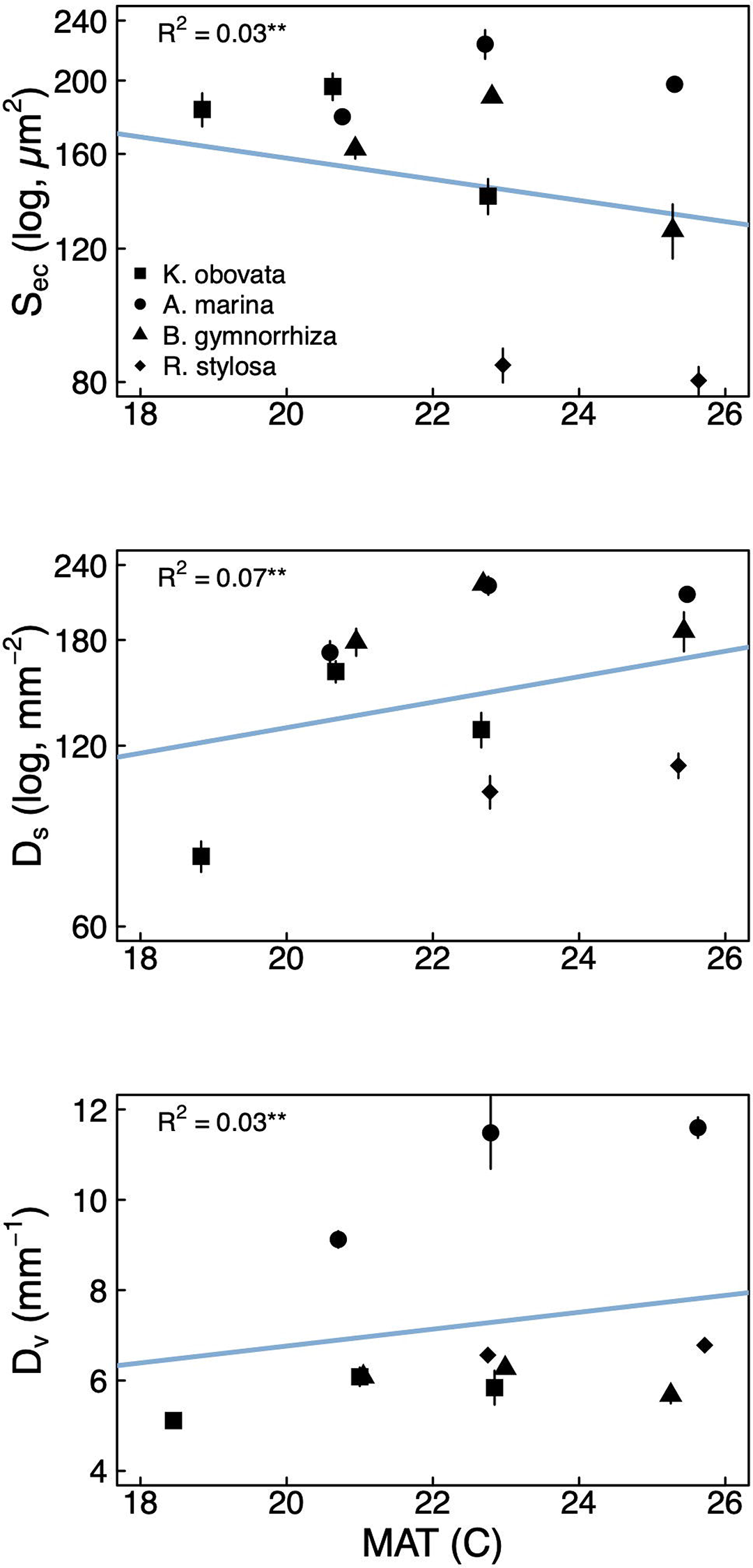
Effects of mean annual temperature (MAT) on leaf anatomical traits of the four species that occurred at multiple sites: (a) epidermal cell size (*S_ec_*), (b) abaxial stomatal density (*D_s_*), and (c) leaf vein density (*D_v_*). Blue lines and the marginal *R^2^* values were calculated by linear mixed effects models based on individual plant level data and indicate the effect of MAT on each trait after accounting for the random effect of species (i.e. each species has a different intercept). For easier visualization, points and error bars represent species x site means and standard error. Note that all traits except vein density (*D_v_*) are plotted on log-scaled y-axes and that slight jitter has been added along the x-axis to better distinguish points. **P < 0.01

### Relationships among leaf anatomical traits: S_s_, D_s_, D_v_, and S_ec_

We compared inter- and intra-specific coordination between epidermal cell size, stomatal size, stomatal density, and vein density to data previously reported for angiosperms (Beaulieu *et al*., 2008; Carins Murphy *et al*., 2012, 2014, 2016) to test for common allometric scaling relationships. The data from Carins Murphy et al. (2012, 2014, 2016) included intraspecific variation driven by growing plants in high and low light or VPD environments. We computed SMA regressions and confidence intervals from the broader selection of angiosperms and compared these to the relationships observed for the 13 mangrove species in this study. In addition, we tested for coordinated trait evolution by calculating generalized least squares regressions among traits that included the expected covariance due to phylogenetic relatedness. Overall, mangroves did not conform to the scaling relationships previously observed for a broader selection of angiosperms (Fig. 4). While most angiosperms had larger epidermal cells than stomata, species with smaller cells overall (i.e. both smaller guard cells and epidermal cells) also had larger stomata than epidermal cells, and all of the mangroves sampled here had epidermal cells smaller than stomata (Fig. 4a). While *S_ec_* strongly and positively scaled with *S_s_* among non-mangrove angiosperms (slope = 0.77 [0.68, 0.88], R^2^ = 0.49, P < 0.001) even after accounting for shared evolutionary history (Fig. S4), the relationship between *S_s_* and *S_ec_* among mangroves was negative (slope = −0.97 [−1.41, −0.66], R^2^ = 0.37, P < 0.01; Fig. 4a) and only marginally significant after accounting for shared evolutionary history (P = 0.068). The relationship between *D_s_* and *S_ec_* was significant and negative among non-mangrove angiosperms (slope = −0.85 [−0.95, −0.76], R^2^ = 0.62, P < 0.001) even after accounting for shared evolutionary history (Fig. S3), but for mangroves the relationship between these traits was positive, though only marginally significant (slope = 0.88 [0.57, 1.36], R^2^ = 0.17, P = 0.07; Fig. 4b) and non-significant after incorporating shared evolutionary history (P = 0.27). Removing the one species (*Laguncularia racemosa*) that had the largest epidermal cells resulted in a significant, positive relationship between *S_ec_* and *D_s_* (slope = 1.00 [0.68, 1.48], R^2^ = 0.40, P < 0.01) that remained significant after accounting for shared evolutionary history (P = 0.03). Because these relationships are fundamentally about packing cells into a two-dimensional space, we used the *D_s_* only of the abaxial leaf surface and not the total *D_s_* per leaf area (i.e. the sum of abaxial and adaxial surfaces, *D_s,tot_*). Including instead the total *D_s_* (sum of adaxial and abaxial) for the three amphistomatous species resulted in a positive, significant relationship between *D_s_* and *S_ec_* (slope = 0.89 [0.61, 1.29], R^2^ = 0.41, P < 0.01) that was marginally significant after accounting for shared evolutionary history (P = 0.06). There was a strong and significant negative relationship between *D_v_* and *S_ec_* for non-mangrove angiosperms (slope = −0.47 [−0.59, −0.38], R^2^ = 0.81, P < 0.001; Fig. 4c), but this relationship was positive though not significant for mangroves either excluding (P = 0.22) or including shared evolutionary history (P = 0.87). Across sites and species, the relationship between stomatal size (*S_s_*) and density (*D_s_*) for mangroves was consistent with the relationship across a broader sampling of angiosperms (Fig. 4d; Fig. S3). Leaves always had smaller stomata or fewer stomata than the maximum theoretical packing limit (solid line in Fig. 4e). Mangrove species with higher *D_s_* had smaller *S*_s_ (non-phylogenetic SMA: R^2^ = 0.42, P < 0.01; phylogenetic GLS: P = 0.03), and the slope of this relationship (slope = −0.91 [−1.32, −0.63]) was not significantly different (P = 0.44) from the slope across all angiosperms (non-phylogenetic: slope = −0.85 [−0.91, −0.79], R^2^ = 0.65, P < 0.0001; phylogenetic: Fig. S3). However, intraspecific variation across sites revealed that the overall negative relationship between *S_s_* and *D_s_* was not always apparent within species (Fig. S4 inset). For example, at intermediate MAP, *Kandelia obovata* had larger *S_s_* and lower *D_s_* than at both low and high MAP, but across all three sites where this species occurred, there was a negative relationship between *D_s_* and *S_s_*. We further tested whether tradeoffs between cell size and packing densities extended to leaf veins (Fig. 4e). While across all angiosperms species with smaller *S_s_* had higher *D_v_* (R^2^ = 0.13, P < 0.0001) even after accounting for shared evolutionary history (Fig. S3), among mangroves there was no significant relationship between *S_s_* and *D_v_* whether (P = 0.54) or not (P = 0.07) shared evolutionary history was included, though the relationship was negative and the mangroves fell within the range of trait values occupied by non-mangrove angiosperms (Fig. 4e).

**Fig. 4.**
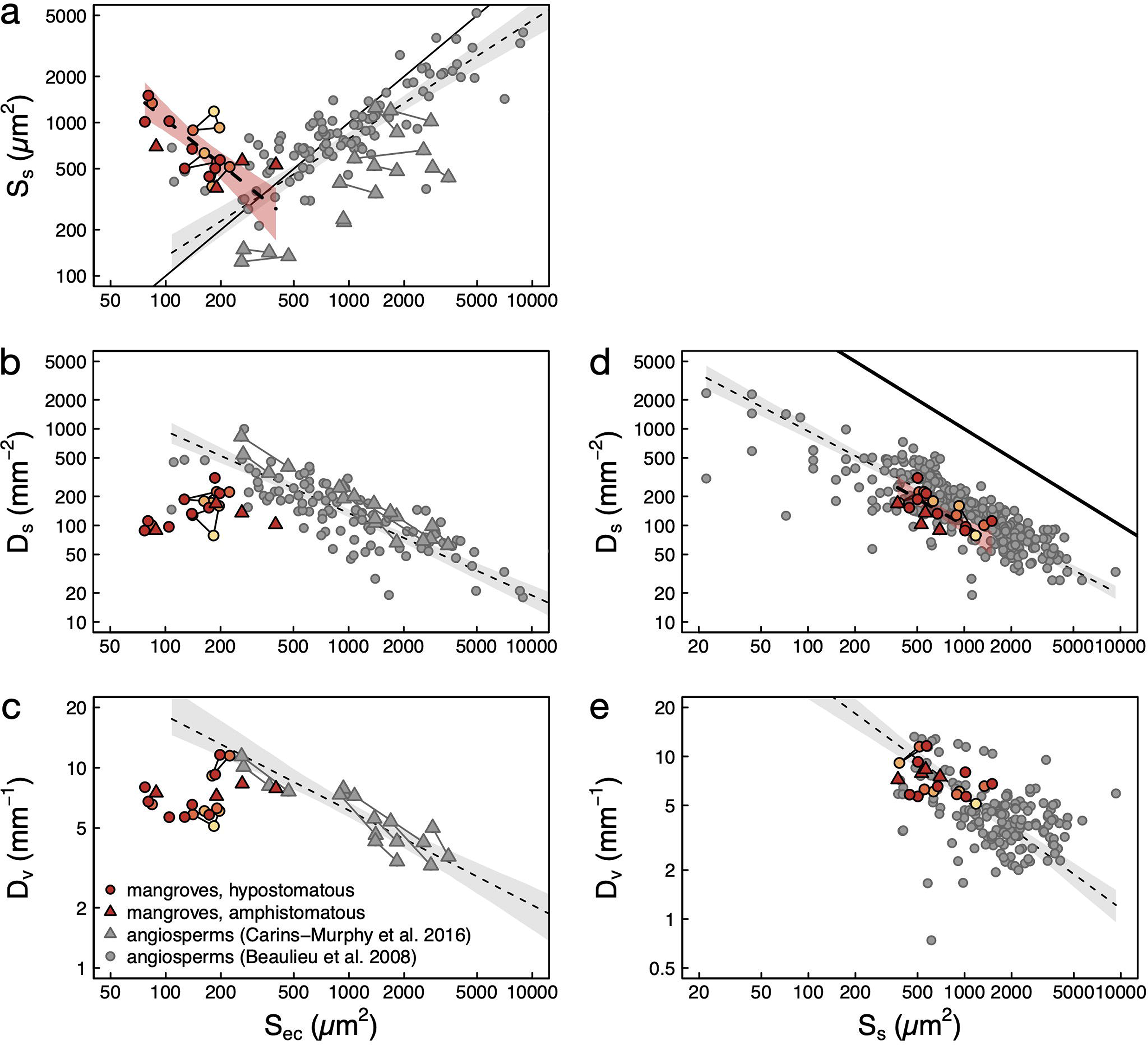
The scaling of epidermal cell size (*S_ec_*) and (a) stomatal size (*S_s_*), (b) stomatal density (*D_s_*), and (c) vein density (*D_v_*), and the scaling of stomatal size (*S_s_*) and (d) stomatal density (*D_s_*) and (e) vein density (*D_v_*). In (a-c), grey circles represent data from Beaulieu et al. (2008), and grey triangles represent data from Carins-Murphy et al. (2016), with lines connecting conspecific plants grown under different conditions. In (a), the solid line represents the 1:1 line. In (d) the thick solid line represents the maximum packing limits where *D_s_* = 1/*S_s_* (Franks and Beerling, 2009). In (d) the inset shows a focused view of mangrove data colored according to site MAP (Figure 1) with solid lines connecting species across multiple sites and dashed lines connecting adaxial and abaxial data for amphistomatous species. In (d-e), grey points are previously published data from a variety of sources (see methods). (b,d) Only *D_s_* of the abaxial (lower) surface is plotted for amphistomatous species because this relationship is based on cell packing. In all panels, grey dashed lines and shading are standard major axis regressions and confidence intervals for angiosperms, and dashed lines and red shading are standard major axis regressions and confidence intervals for mangroves.

### Coordination between D_v_, D_s, tot_ and maximum theoretical stomatal conductance

Although *D_s,tot_* and *D_v_* were coordinated among all the mangrove species x site combinations sampled here (slope = 1.63 [1.08, 2.45], R^2^ = 0.28, P = 0.02), *D_s,tot_* was generally lower for a given *D_v_* than it was among a broader sampling of angiosperms (slope = 1.53 [1.37, 1.71], R^2^ = 0.37, P < 0.001; Fig. 5a). For non-mangrove angiosperms, the relationship between *D_s,tot_* and *D_v_* remained as strong after accounting for shared evolutionary history (Fig. S5), but for mangroves there was no coordinated evolution of *D_s,tot_* and *D_v_* (P = 0.84). Mangroves maintained a similar maximum theoretical stomatal conductance for a given *D_s,tot_* as other non-mangrove angiosperms (mangrove slope = 0.56 [0.43, 0.75], R^2^ = 0.68, P < 0.0001, angiosperm slope = 0.64 [0.60, 0.68], R^2^ = 0.68, P < 0.0001, slope test P = 0.39; Fig. 5b), and there was evidence for coordinated evolution between *D_s,tot_* and maximum theoretical *g_s_* for both mangroves (P < 0.01) and non-mangrove angiosperms (Fig. S5).

**Fig. 5.**
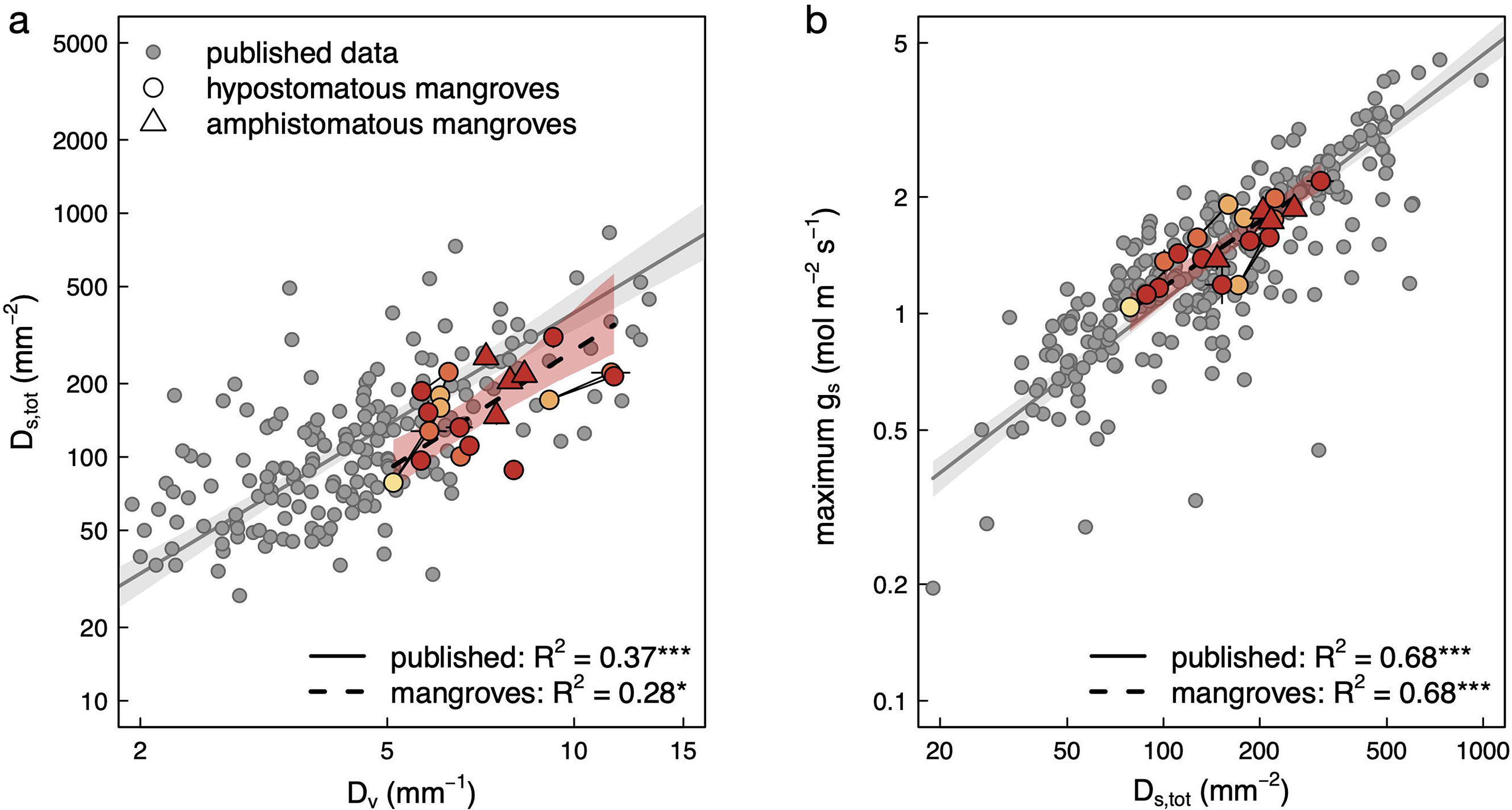
Coordination between (a) vein density (*D_v_*) and total stomatal density (*D_s,tot_*, the sum of adaxial and abaxial *D_s_*, relevant for fluxes) and (b) total stomatal density (*D_s,tot_*) and maximum stomatal conductance (*g_s_*) for mangroves (yellow/red points, colored according to site MAT, and connected by solid lines for species occurring in multiple sites) compared to a broader sampling of angiosperms (grey points). Circles represent species with hypostomatous leaves, and triangles represent species with amphistomatous leaves. The solid, grey lines and shading are the SMA regressions and 95% CI for angiosperms, and the dashed, black lines and red shading are the SMA regressions and 95% CI for mangroves.

### Relationships between leaf size, LMA, S_ec_, and D_ec_

We also tested how these anatomical traits may be related to intra- and interspecific variation in leaf size, which varied approximately five-fold across the mangrove species sampled here. Across species, larger leaves were significantly associated with higher LMA (R^2^ = 0.23, P = 0.02; Fig. 6a), smaller epidermal cells (R^2^ = 0.20, P < 0.05; Fig. 6b), and a higher packing density of epidermal cells (R^2^ = 0.46, P < 0.001; Fig. 6c). The relationship between leaf size and epidermal cell size was even stronger when the one mangrove species with very large epidermal cells (*Laguncularia racemosa*) was removed (R^2^ = 0.27, P < 0.05). However, these pairwise relationships were weaker or not significant after accounting for shared evolutionary history: there was no significant relationship between LMA and leaf size (P = 0.34), there was no significant relationship between *S_ec_* and leaf size (P = 0.25) though excluding *L. racemosa* improved the relationship (P = 0.065), and there was a marginally significant relationship between *D_ec_* and leaf size (P = 0.073).

**Fig. 6.**
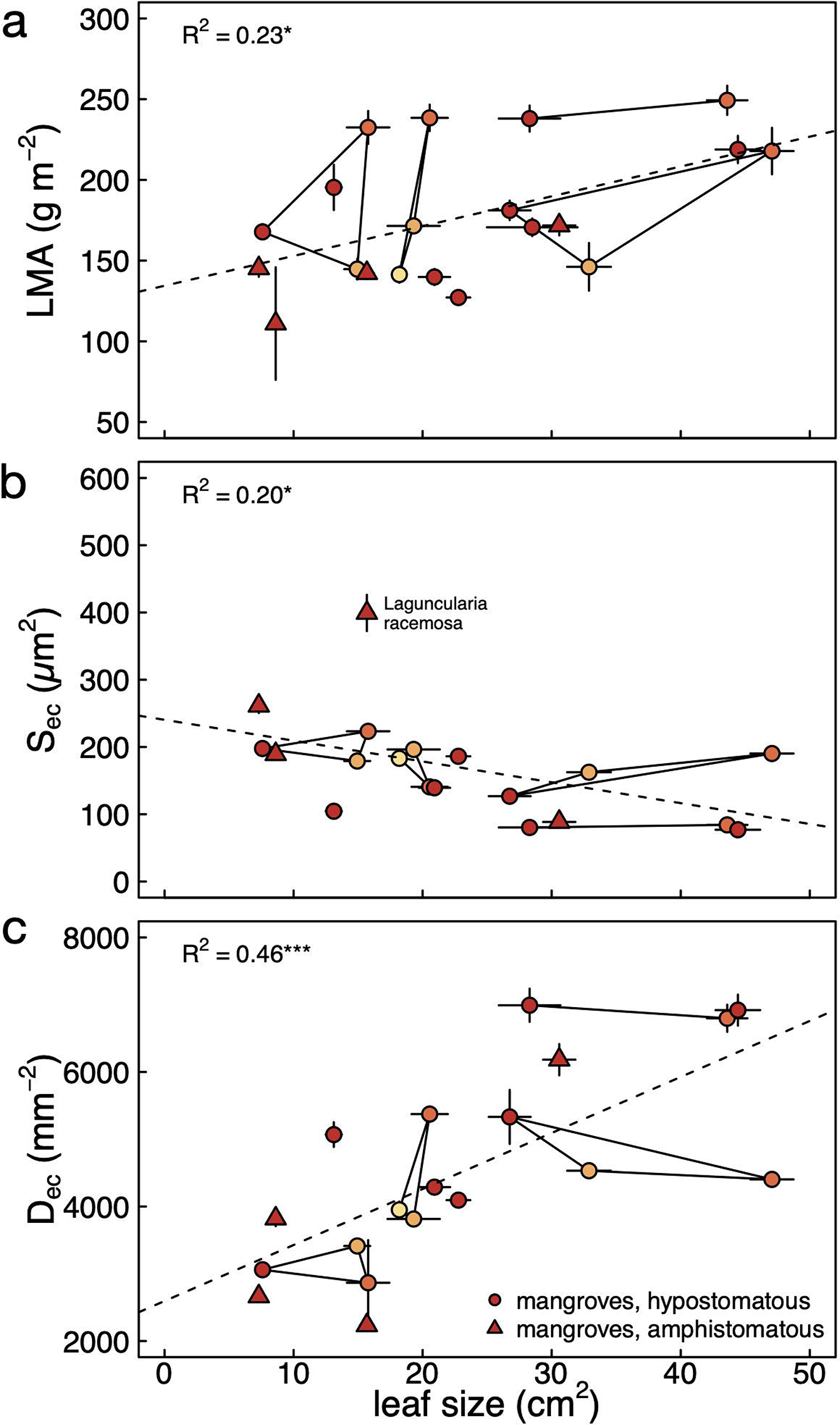
Relationships between leaf size and (a) leaf mass per area (LMA), (b) epidermal cell size (*S_ec_*), and (c) packing density of epidermal cells (*D_ec_*). Connected points represent species that occur across multiple sites, and points are colored according to site-specific MAT.

## DISCUSSION

Our analysis of 13 mangrove species, four of which occurred at more than one site, provides strong evidence that the allometry of cells and tissues in mangrove leaves is distinct from other C_3_ angiosperm species. Our results highlight that while mangroves exhibit some of the same trait relationships exhibited by non-mangrove angiosperms, they deviate in some potentially important ways, most notably that they have unusually small epidermal pavement cells and large guard cells. Despite these deviations from other angiosperms, mangroves nonetheless attained similar maximum theoretical stomatal conductance. Because leaves are composed of multiple cell types and because genome size limits only minimum cell size, there can be numerous combinations of final cell sizes and packing densities that allow for variation in leaf structure that lead to similar maximum potential gas exchange. Understanding the implications of these differences could further shed light on how the unique selective pressures of the mangrove habitat have resulted in novel anatomical and physiological adaptations.

The mangroves we sampled from four sites in China (Fig. 1) had relatively small genomes compared to other terrestrial vascular plants, yet they did not necessarily have smaller cells. While interspecific analyses of genome size and cell size have repeatedly shown positive scaling between genome size and minimum cell size, the absolute range of cell sizes is negatively related to genome size, with smaller genomes allowing for a greater range of final cell sizes (Beaulieu *et al*., 2008; Simonin and Roddy, 2018; Roddy *et al*., 2020; Théroux-Rancourt *et al*., 2021). The mangroves sampled here highlight this important nuance in the genome size-cell size relationship: smaller genomes allow for smaller cells, but smaller genomes do not necessarily mean that cells will always be small (Simonin and Roddy, 2018; Roddy *et al*., 2020). The interspecific relationship between stomatal guard cell size and genome size among mangroves was actually negative, i.e. species with larger genomes had smaller guard cells (Fig. 2). It is important to note that this negative relationship does not contradict previous analyses (e.g. Beaulieu *et al*., 2008) because a random sampling of any subset of species in these previous analyses–particularly a set of species that exhibits little variation in genome size–could produce the same negative relationship. Furthermore, the inter- and intraspecific variation in guard cell size reported here all occurred within the range of sizes defined by previously published data (Fig. 2, 4), and all of these guard cells were larger than the minimum cell volume modeled from genome size. Similarly, while epidermal cells were smaller than guard cells in all species, they were always larger than the minimum cell size modeled from genome size (Fig. 2), further reiterating that genome size is associated with strict limits on minimum cell size but has less direct impact on maximum or mature cell sizes (Roddy *et al*., 2020).

Above the minimum cell size defined by the size of the genome, leaf cell sizes and packing densities can vary in response to abiotic conditions (Blonder *et al*., 2017, Veselý *et al*., 2020, Wang *et al*., 2020, Zhao *et al*., 2020). Using the four species that occurred in more than one site (Table 2), we calculated the effects of MAT, MAP, and soil water salinity on each anatomical trait (Fig. 3 and S2, Table S1). Interestingly, MAT and soil water salinity, which were strongly correlated among the four sites, affected most traits, whereas MAP had no effect. Overall these environmental effects were weak, with no single environmental variable explaining more than 8% of the variation in a trait, similar to the relatively weak effects on leaf traits of mangrove seedlings grown under different temperatures (Inoue *et al*., 2021). Some of the significant effects of MAT were driven by the coldest site (FD), where only one species (*Kandelia obovata*) was present (Fig. 1, Table 2). Although these trait correlations with climate showcase that cell and tissue traits are plastic and that this plasticity is limited by the minimum cell size defined by genome size, the climate effects were relatively weak with differences among species explaining a greater proportion of the trait variance (Table S1).

While there is typically a tradeoff between stomatal size and density (Franks and Beerling, 2009) that is driven largely by genome size variation among species (Beaulieu *et al*., 2008; Simonin and Roddy, 2018; Roddy *et al*., 2020; Veselý *et al*., 2020; Théroux-Rancourt *et al*., 2021), individual species can move through this bivariate phenotype space in different ways (Fig. S4). Although *S_s_* and *D_s_* for the 13 mangrove species showed a negative relationship that overlapped with previous observations of angiosperms (Fig. 4d, Fig. S4), a strict, mechanistic tradeoff between *S_s_* and *D_s_* occurs only at the packing limit (solid line in Fig. 4d), and as species move farther away from this packing limit the potential for a strict tradeoff between size and density becomes less likely. The species-specific responses to MAT highlight that because both *S_s_* and *D_s_* are far from their packing limit (solid line in Fig. 4), a relatively wide range of *D_s_* can occur for a given *S_s_*. For example, in *Kandelia obovata*, decreasing temperature from the warmest site causes an increase in *D_s_* with almost no change in *S_s_*, but decreasing MAT to the coolest site causes a reduction in *D_s_* and a large increase in *S_s_* (Fig. S4 inset). These intraspecific patterns also highlight that traits do not necessarily covary within species across habitats the same way they do among species. Furthermore, while genome size determines minimum cell sizes and maximum cell packing densities (Fig. 2), acclimation and adaptation of mature cell sizes and cell packing densities can be driven by the environment independent of genome size (Jordan *et al*., 2015) or vary due to other species-specific traits or constraints.

Despite the cell type-specific and species-specific responses of leaf anatomy to environmental conditions (Figs. 3, 4, S2), *D_s,tot_* and *D_v_* were strongly and positively related (Fig. 5a)–though not after accounting for shared evolutionary history–and mangroves attained the same maximum theoretical *g_s_* for a given *D_s_* as other angiosperms (Fig. 5b). The coordination between *D_s,tot_* and *D_v_* across environments occurred among species and, generally, within species, although there was variation among species in the intraspecific trends (Figs. 4, 5). The coordination between *D_s_* and *D_v_* within and across species has been attributed to changes in the size and density of epidermal pavement cells (Carins Murphy *et al*., 2012, 2014, 2016, 2017). Specifically, epidermal cell size is thought to depend on environmental conditions, such that differential expansion of epidermal cells modulates the spacing of stomata and bundle sheath extensions from the veins. For the angiosperms studied so far, inter- and intraspecific coordination between epidermal cell size (*S_ec_*) and both *D_s_* and *D_v_* have been taken as evidence in support of this ‘passive dilution’ model (Carins Murphy *et al*., 2012, 2014, 2016). In contrast to sun and shade leaves of nine angiosperms species (Carins Murphy *et al*., 2012) and a broader sampling of angiosperms (Beaulieu *et al*., 2008), mangroves deviated in the relationships between *S_ec_* and both *D_s_* and *S_s_* (Fig. 4). While larger epidermal cells are typically associated with larger stomata, among mangroves this relationship was negative, due at least partially to the fact that mangrove epidermal cells were substantially smaller than their stomata (Figs. 1, 2, 4a). The one mangrove species that had epidermal cells larger than stomata was *Laguncularia racemosa*, and excluding this one species revealed a significant, positive relationship between *S_ec_* and *D_s_*, in contrast to the negative relationship between *S_ec_* and *D_s_* reported from other angiosperms (Fig. 4b). Additionally, there was no relationship between *S_ec_* and *D_v_* in mangroves, in contrast to the negative inter- and intraspecific relationship previously reported for nine non-mangrove angiosperm species (Fig. 4c). Contrary to the lack of correlation between *S_ec_* and both *D_s_* and *D_v_*, the 13 mangroves studied here overlap with the broader group of angiosperms showing a negative correlation between *S_s_* and both *D_s_* and *D_v_* (Figs. 4d and 4e). Therefore, while there is coordination between *D_v_* and *D_s_* among mangroves (Fig. 5a), variation in epidermal cell size is likely not responsible for maintaining this coordination either within species across environmental conditions or among species. That only some of these leaf traits exhibited correlated evolution among mangroves could be due to the relatively small sample size of only 13 species and to the relatively small variation in traits exhibited by mangroves compared to the full range of trait values exhibited by non-mangrove angiosperms. Nonetheless, these patterns in mangrove anatomy suggest that the unusually small epidermal cells and the variation in *S_ec_* within and between species may influence other aspects of leaf function important to the mangrove habit, such as osmotic balance or leaf biomechanics.

All else being equal, smaller cells are more resistant to mechanical buckling than larger cells (Terashima *et al*., 2001). Additionally, cellular biomechanics is intimately related to cell water balance via the effects of wall thickness on the bulk elastic modulus and the sensitivity of turgor pressure to changes in water content. Possessing small and numerous epidermal cells may be particularly advantageous for plants living in saline conditions that impose an osmotic stress on cells throughout the plant. Indeed, *S_ec_* decreased with increasing temperature and salinity (Figs. 3 and S2), as would be predicted if osmotic balance and cell mechanics were linked to epidermal cell size. Compared to freshwater and coastal plants, marine plants have much stiffer cell walls that allow them to maintain the high turgor pressures necessary to tolerate low osmotic potentials (Touchette *et al*., 2014). While we do not have water potential data or pressure-volume curve parameters for the mangroves studied here, previous studies suggest that mangroves usually have lower water potentials than other terrestrial angiosperms (Jiang *et al*., 2017, 2021, 2022). Based on these lines of evidence, we predict that lower osmotic potentials would be associated with hotter, more saline conditions and would be related to epidermal cell size. Further evidence that epidermal cells may be important for the biomechanics of mangrove leaves comes from the strong–and unexpected– relationships between *S_ec_* and *D_ec_* and leaf size (Fig. 6). Larger leaves, which also have higher LMA, have smaller and more densely packed epidermal cells (Fig. 6). However, there were no significant relationships between leaf size and either *S_s_* or *D_s_* (data not shown), in contrast to relationships seen in *Rhizophora mangle* across salinity gradients (Peel *et al*., 2017). In addition to being small, epidermal cells in mangrove leaves were also more circular (Fig. 1) than epidermal cells of most other angiosperms, which are often highly invaginated and puzzle-shaped (Vofely *et al*., 2019). Puzzle-shaped cells seem to develop in order to reduce mechanical stress without requiring excessively thick walls (Sapala *et al*., 2018). The small, densely packed epidermal cells in mangroves may be advantageous because they increase mechanical stiffness, allowing for larger leaves.

The warm, windy, saline environments of the mangrove habitat have driven the evolution of a variety of physiological strategies, including tolerance to low osmotic potentials, salt exclusion, and salt secretion, all of which influence mangrove hydraulics and photosynthesis (Ball, 1988; Sobrado, 2000, 2002; Jiang *et al*., 2017, 2022). Mangroves had higher *D_v_* for a given *D_s_* than non-mangrove angiosperms, yet their environment and physiology are amenable to foliar water uptake, which is expected to relax selection for high vein densities. Deliquescence of salts on mangrove leaves can facilitate foliar water uptake even when atmospheric humidity is unsaturated (Coopman *et al*., 2021). The mean relative humidities at the four sites where we sampled were all within the range in which deliquescence of salts is likely (Zeng *et al*., 2013). Thus, foliar water uptake may play an important role in mangrove leaf water balance (Schreel *et al*., 2019), potentially relaxing the role of leaf venation in efficiently providing all of the water to the leaf. The likelihood that mangrove leaves may use both root-derived and atmospheric water to hydrate leaves may relax selection on the xylem to efficiently supply water and result in greater variation in leaf structure-function relationships. Furthermore, that mangroves had smaller epidermal cells than their stomata, in contrast to non-mangrove angiosperms, highlights another adaptation of mangrove anatomy that may be advantageous in the warm, saline mangrove habitat. Understanding the implications of small epidermal cells on mangrove hydraulics, gas exchange, and biomechanics would be an important advance in understanding mangrove leaf adaptations.

## CONCLUSION

Our results show that mangroves attain similar maximum theoretical gas exchange capacity to other angiosperms despite deviating in many anatomical relationships well-characterized for angiosperms. This highlights that there are multiple anatomical solutions with the same functional outcome. The small genomes of mangroves allow for large variation in cell sizes and cell packing densities in response to abiotic conditions. The unusually small epidermal cells of mangrove leaves may help them tolerate the mechanical and osmotic demands of their saline environments and their leaf size. Whether the extremely small epidermal cells that enhance cell packing is an adaptation to the stressful mangrove environment deserves further investigation.

## SUPPLEMENTARY DATA

Additional Supporting Information may be found in the online version of the article at the publisher’s website.

**Table S1**: Linear mixed effects model results of each environmental parameter (MAP, soil salinity, MAT) on each anatomical trait for the four species that occurred at multiple sites.

**Figure S1**: Phylogenetic regression statistics (slope, *t*, *P*) of non-mangrove angiosperms for relationships presented in Figure 2.

**Figure S2**: The effects of mean annual precipitation (MAP), soil salinity, and mean annual temperature (MAT) on leaf anatomical traits of the four species that occurred at multiple sites.

**Figure S3**: Phylogenetic regression statistics (slope, *t*, *P*) of non-mangrove angiosperms for relationships presented in Figure 4.

**Figure S4**: The relationship between stomatal size (*S_s_*) and stomatal density (*D_s_*) for non-mangrove angiosperms (grey points) and the mangrove species samples here (yellow-red points).

**Figure S5**: Phylogenetic regression statistics (slope, *t*, *P*) of non-mangrove angiosperms for relationships presented in Figure 5.

## FUNDING

This work was supported by grants from the National Natural Science Foundation of China (grant number 31860195) and Natural Science Foundation of Guangxi (Key Program 2022GXNSFDA035059) to G-FJ, and by DEB-1838327 from the U.S. National Science Foundation to KAS and ABR.

## ACKNOWLEDGMENTS

We thank Guangxi Mangrove Research Center, Guangxi Academy of Sciences for providing soil water salinity data, and the many people (especially De-Hua Zeng from Sanya Academy of Forestry) who helped with field sampling. This manuscript benefitted from constructive comments by two reviewers. The authors declare no conflict of interest.

## AUTHORS’ CONTRIBUTIONS

G.-F.J. conceived the ideas and designed the study. G.-F.J., S.-Y. Li, and A.B.R. collected the data. G.-F.J. and A.B.R. analyzed the data. G.-F.J., A.B.R. and K.A.S. wrote the manuscript, and all authors reviewed each draft before giving approval for submission of the final version.

---

**Figure S1**. The effects of mean annual precipitation (MAP), soil salinity, and mean annual temperature (MAT) on leaf anatomical traits of the four species that occurred at multiple sites: (a,f,k) stomatal size (*S_s_*), (b,g,l) epidermal cell size (*S_ec_*), (c,h,m) abaxial stomatal density (*D_s_*), (d,i,n) epidermal cell packing density (*D_ec_*), and (e,j,o) leaf vein density (*D_v_*). Blue lines indicate the effect of each environmental variable on each trait after accounting for the random effect of species (i.e. each species has a different intercept). Note that all traits except vein density (*D_v_*) are plotted on log-scaled y-axes. Points represent individual plants, whereas in Figure 3 points represent species x site means. See Table S1 for complete marginal and conditional *R^2^* values.

**Table S1**. Linear mixed effects model results of each environmental parameter (MAP, soil salinity, MAT) on each anatomical trait for the four species that occurred at multiple sites. Note that the marginal *R^2^* is the proportion of the variance explained by the fixed effects alone (i.e. the environmental variable) and that the conditional *R^2^* is the proportion of the variance explained by both the fixed and random effects (i.e. the environmental variable and the species identity). *P < 0.05; **P < 0.01, ***P < 0.001

## Notes

### Competing Interest Statement

The authors have declared no competing interest.

