## Supplemental Information for "Mangroves deviate from other angiosperms in their genome size, leaf cell size, and cell packing density relationships"

**Table S1.** Linear mixed effects model results of each environmental parameter (MAP, soil salinity, MAT) on each anatomical trait for the four species that occurred at multiple sites. Note that the marginal *R^2^* is the proportion of the variance explained by the fixed effects alone (i.e. the environmental variable) and that the conditional *R^2^* is the proportion of the variance explained by both the fixed and random effects (i.e. the environmental variable and the species identity). *P < 0.05; **P < 0.01, ***P < 0.001

| trait | environmental variable | marginal *R^2^* | conditional *R^2^* |
| --- | --- | --- | --- |
| stomatal size (*S_s_*) | MAP | 0 | 0.90 |
|  | salinity | 0 | 0.90 |
|  | MAT | 0 | 0.90 |
| epidermal cell size (*S_ec_*) | MAP | 0 | 0.86 |
|  | salinity | 0.02 | 0.85 |
|  | MAT | 0.03** | 0.86 |
| stomatal density (*D_s_*) | MAP | 0 | 0.72 |
|  | salinity | 0.08*** | 0.80 |
|  | MAT | 0.07** | 0.78 |
| epidermal cell density (*D_ec_*) | MAP | 0 | 0.54 |
|  | salinity | 0.01 | 0.53 |
|  | MAT | 0.01 | 0.53 |
| vein density (*D_v_*) | MAP | 0 | 0.87 |
|  | salinity | 0.03** | 0.89 |
|  | MAT | 0.03** | 0.88 |

**Figure S1.** Phylogenetic regression statistics (slope, *t*, *P*) of non-mangrove angiosperms for relationships presented in Figure 2. Distributions of phylogenetic regression statistics are based on 1000 equally likely phylogenies. Vertical red lines indicate where *P* = 0.05. (top row): Statistics for the relationship between guard cell size (Sgc) and genome size. (bottom row): Statistics for the relationship between epidermal cell size (Sec) and genome size.

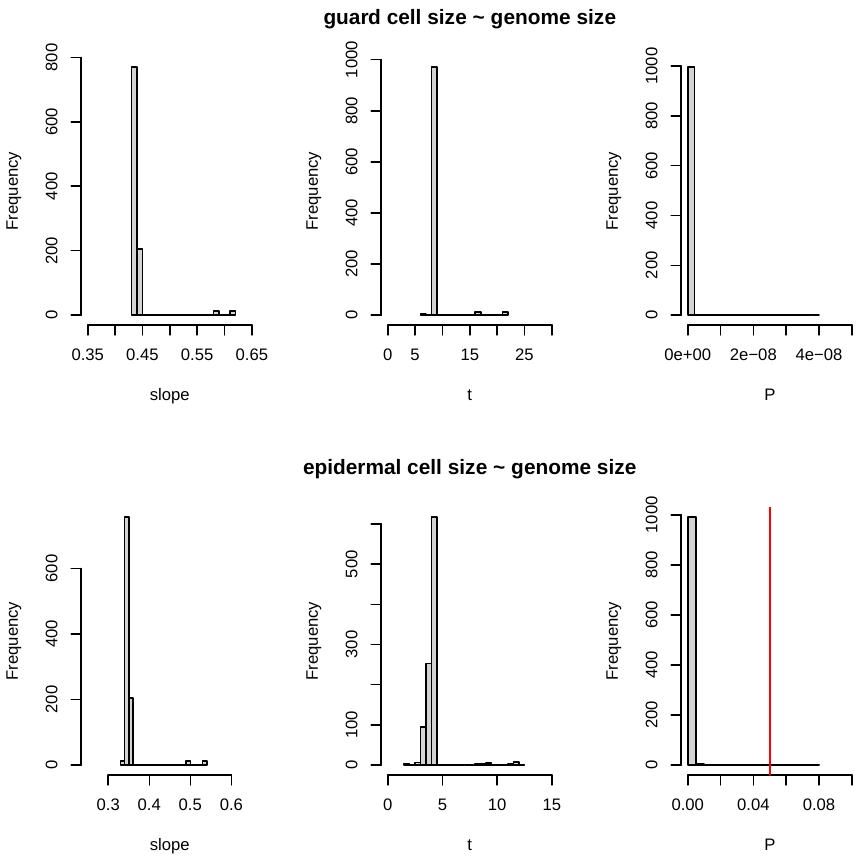

**Figure S2.** The effects of mean annual precipitation (MAP), soil salinity, and mean annual temperature (MAT) on leaf anatomical traits of the four species that occurred at multiple sites: (a,f,k) stomatal size (*S_s_*), (b,g,l) epidermal cell size (*S_ec_*), (c,h,m) abaxial stomatal density (*D_s_*), (d,i,n) epidermal cell packing density (*D_ec_*), and (e,j,o) leaf vein density (*D_v_*). Blue lines indicate the effect of each environmental variable on each trait after accounting for the random effect of species (i.e. each species has a different intercept). Note that all traits except vein density (*D_v_*) are plotted on log-scaled y-axes. Points represent individual plants, whereas in Figure 3 points represent species x site means. See Table S1 for complete marginal and conditional *R^2^* values.

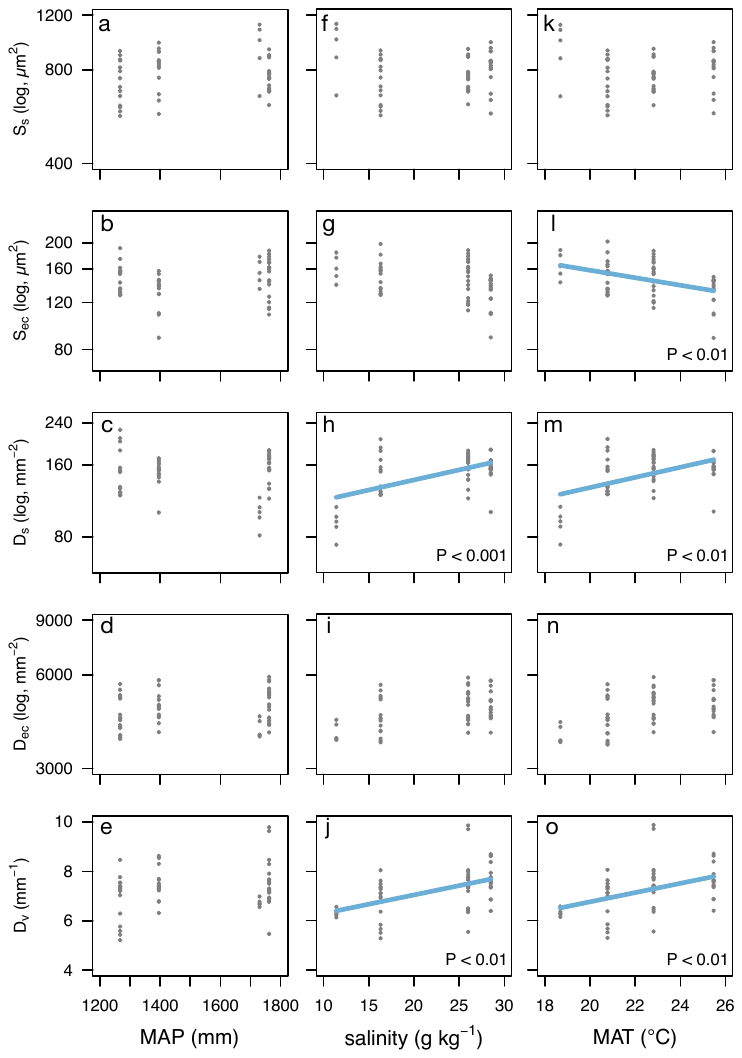

**Figure S3.** Phylogenetic regression statistics (slope, *t*, *P*) of non-mangrove angiosperms for relationships presented in Figure 4. Distributions of phylogenetic regression statistics are based on 1000 equally likely phylogenies. Vertical red lines indicate where *P* = 0.05.

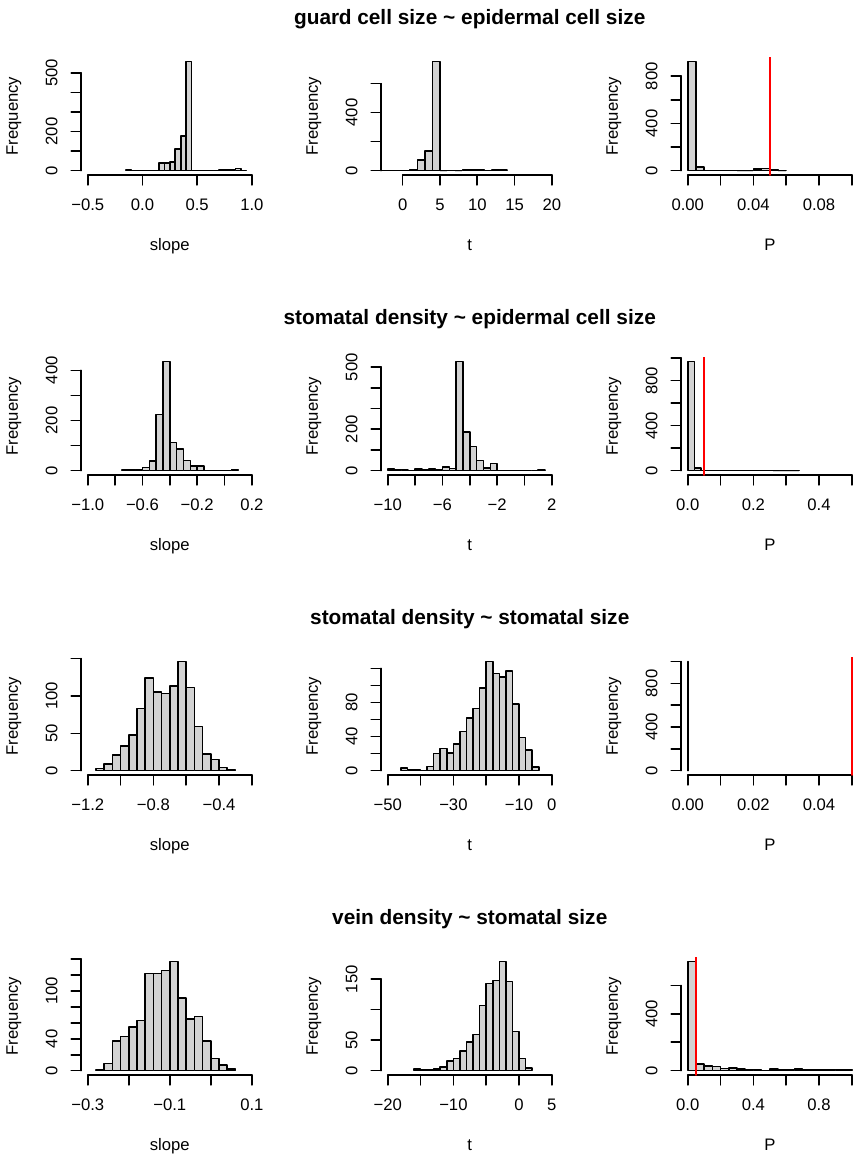

**Figure S4.** The relationship between stomatal size (*S_s_*) and stomatal density (*D_s_*) for non-mangrove angiosperms (grey points) and the mangrove species samples here (yellow-red points). The inset shows only the mangrove species. Points for mangroves are colored according to MAT (see Figure 1). Triangles represent adaxial and abaxial stomata for species that are amphistomatous, and points connected by lines represent the same species that occurs at different sites. The solid line in the main figure is the biophysical packing limit ($D_{s}=S_{s}^{-1}$), according to Franks and Beerling (2009).

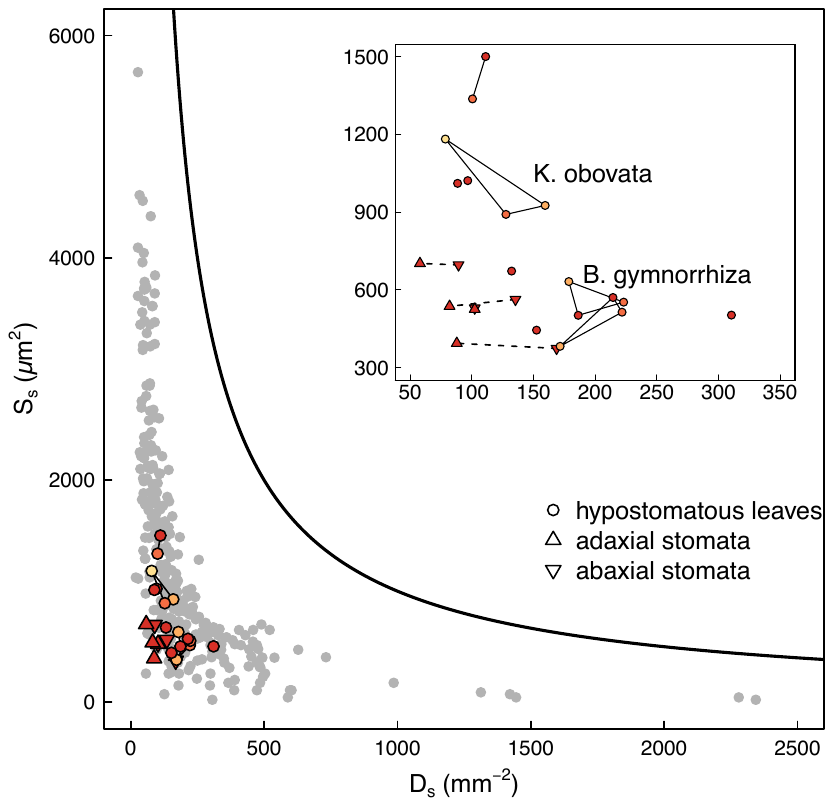

**Figure S5.** Phylogenetic regression statistics (slope, *t*, *P*) of non-mangrove angiosperms for relationships presented in Figure 5. Distributions of phylogenetic regression statistics are based on 1000 equally likely phylogenies. Vertical red lines indicate where *P* = 0.05.

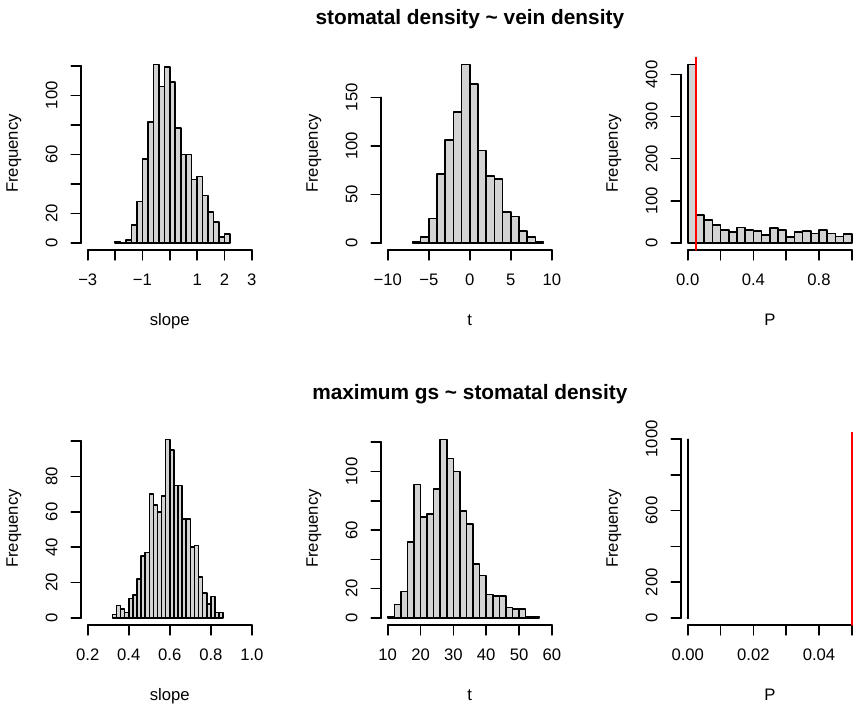
